# DanioDecima: A DNA sequence-to-function model of zebrafish embryogenesis

**DOI:** 10.64898/2026.05.29.728876

**Authors:** Mathias J. Voges, Yang Joon Kim, Max Frank, Benjamin Iovino, Yasin Senbabaoglu, Loic A. Royer

## Abstract

Deep learning DNA sequence-to-function models offer the promise of gaining mechanistic insights into genome regulation, however their performance is often limited by data scarcity in the species of interest. We present DanioDecima, a zebrafish-specific model leveraging transfer learning from human and mouse-trained models to predict tissue- and cell-type-specific gene expression during zebrafish embryogenesis. Initializing DanioDecima with pretrained human and mouse Borzoi and Decima weights raises the median pseudobulk Pearson *r* substantially across cell-types and improves gene-level correlations of test set genes. An *in silico* directed-evolution loop guided by DanioDecima scoring generated synthetic promoters whose motif architectures cluster by the expected target lineage. These findings exemplify a cross-species transfer learning methodology for sequence-to-function models, and position DanioDecima as a practical resource for zebrafish regulatory engineering.

## 1 Introduction

The accurate prediction of genome function from DNA sequence, referred to as the sequence-to-function or regulatory modeling problem, remains a central challenge in genomics. Recent work has emphasized the potential for deep learning models that predict molecular phenotypes from DNA sequence to learn grounded representations of regulatory genomics in human and mouse, as evidenced by the development of models like Enformer, Borzoi, and AlphaGenome [1, 2, 3]. In particular, the use of deep convolutional neural networks (CNNs) and transformer hybrid architectures alongside large-scale functional genomics (e.g., ENCODE [4]), has enabled state-of-the-art models capable of predicting a range of tissue and cell-type functional genomics assays in a multi-task setup [5, 6, 7].

Despite growing capabilities for models in data-replete species, the performance of species-specific bespoke models remains dependent on the quality and kind of functional genomics data available. There have been notable demonstrations of cross-species generalization of a sequence-to-function model between human and mouse [8] – both species with similarly rich data properties – yet performance in other vertebrate species remains underexplored.

Parallel advances in cross-species machine learning are enabled by the emergence of genome-scale foundation models, many of which are pretrained on genomes spanning the evolutionary tree of life, and highlight the propensity of these models to learn representations of the DNA regulatory code in a self-supervised fashion [9, 10, 11]. These DNA language models serve as valuable priors in variant effect prediction [12], and for downstream prediction tasks after fine-tuning, such as gene expression prediction across tissues and cell lines of a given species [10, 13]. While DNA foundation models set a precedent for genome-based cross-species transfer learning, this transfer does not fully benefit from the functional genomics data specifically acquired in these species. Furthermore, DNA language models, fine tuned for a specific functional genomics task, show inconsistencies in their representational awareness of the true regulatory structure [14], likely necessitating the development of foundation models that utilize rich functional genomic priors in addition to solely genomes.

The generalization of sequence-to-function prediction beyond human and mouse is often limited by the paucity of species-specific functional data. Research into zebrafish (*Danio rerio*), an important vertebrate model system for developmental biology and disease, has recently benefited from the generation of a cell type–resolved single-cell atlas of embryogenesis [15]. With this dataset, zebrafish fits within a typical range of data available for a model organism.. Utilizing this dataset, we explore a transfer learning approach as a promising strategy to bridge the gap in model performance between data-rich and data-sparse species, here for a vertebrate system that diverged from mammals over 450 million years ago.

We present DanioDecima, a sequence-to-expression deep learning model for *Danio rerio* embryogenesis. DanioDecima leverages a “pretrain on abundance, fine-tune on scarcity” strategy – initializing with published Borzoi and Decima weights from human and mouse, before adapting to stage- and cell type–resolved zebrafish expression profiles (Fig. 1). This approach improves gene expression prediction relative to random initializations, while also enabling biological interpretability through motif discovery and supporting in silico regulatory design. Together, these capabalities position DanioDecima as a practical framework for engineering zebrafish regulatory elements. Our work demonstrates how the convergence of cross-species datasets, transfer learning, and deep sequence models can aid in the development of predictive models for data-scarce organisms.

**Figure 1:**
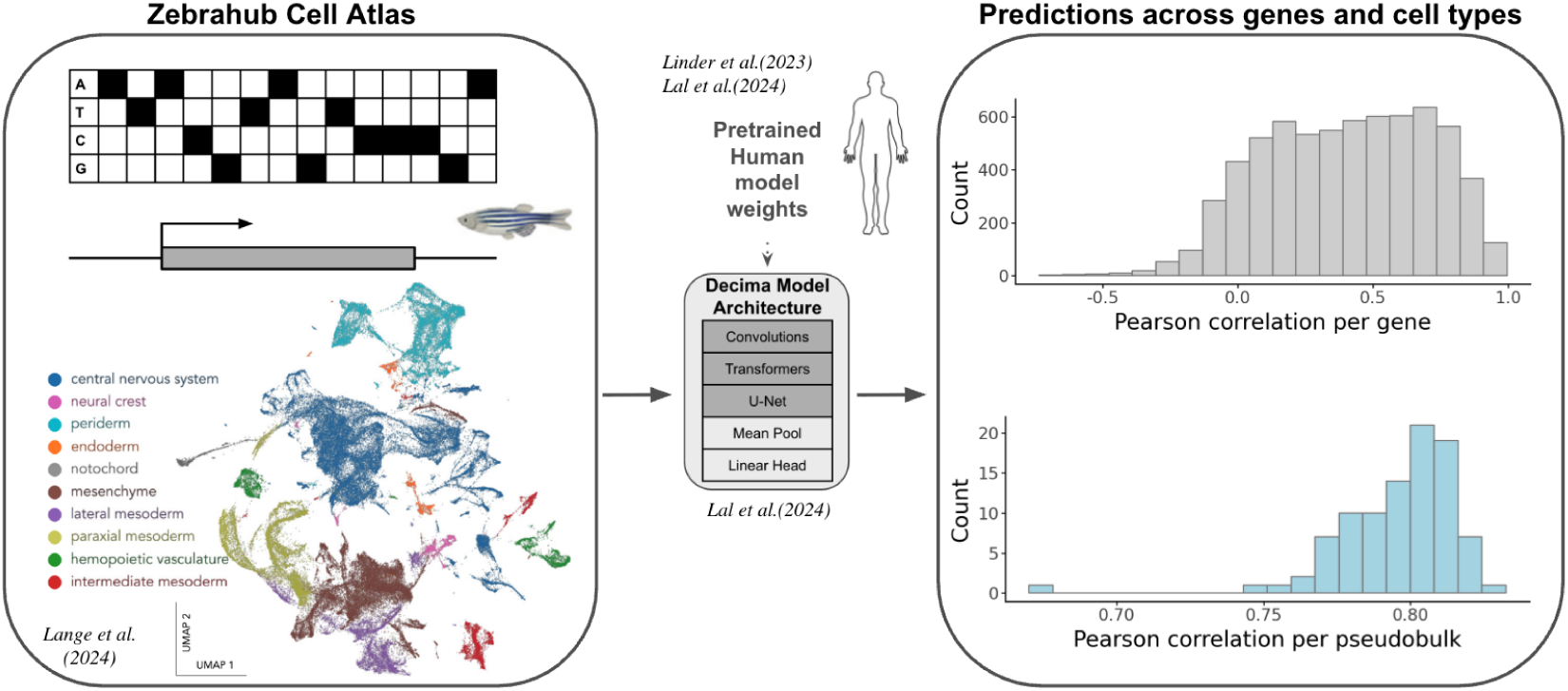
DanioDecima is a sequence-to-expression deep learning model for *Danio rerio* embryogenesis. DanioDecima is initialized with published Borzoi and Decima weights from human and mouse, prior to fine-tuning with stage- and cell type–resolved zebrafish expression profiles.

## 2 Background and Related Work

While prior efforts such as DeepArk have trained deep neural networks to predict cis-regulatory activities (e.g. chromatin marks, TF-bound regions) in *Danio rerio* using DNA sequence alone [16], these approaches primarily focused on regulatory activity prediction rather than gene-level RNA expression or cross-species transfer learning from mammalian pretrained models. Similarly, the Nvwa framework predicts single-cell gene expression landscapes across zebrafish, fruit fly, and earthworm using multi-species single-cell atlases [17]. In contrast, our approach investigates whether large mammalian sequence-to-function models can be transferred to zebrafish developmental gene regulation and applied to regulatory sequence design.

Recent mammalian sequence-to-function models, such as Borzoi and Decima, provide a strong foundation for cross-species regulatory modeling by learning long-range cis-regulatory representations directly from DNA sequence. They have achieved strong predictive performance in human and mouse by training on extensive functional genomics data across many tissues and cell types [2, 7]. Because many aspects of cis-regulatory logic, including transcription factor binding preferences, enhancer-promoter interactions, and sequence determinants of gene expression, are evolutionarily conserved, these models are likely to learn regulatory representations that generalize beyond the human and mouse datasets on which they were trained.

DanioDecima is, to our knowledge, the first model to leverage cross-species transfer learning from pretrained human/mouse regulatory models (Borzoi, Decima) to fine-tune on cell-type–resolved zebrafish expression across embryonic development; and to reveal cell-level mechanistic insights of sequence regulatory grammar in zebrafish via the design of synthetic regulatory elements through directed evolution. This combination of cross-species initialization, developmental time-series fine-tuning, and *in silico* synthetic regulatory element design distinguishes DanioDecima and represents a valuable resource for modeling regulatory genomics and the accurate regulatory DNA sequence design in model organisms with sparse data.

## 3 Methods

DanioDecima extends the Decima modeling framework [7] for cross-species regulatory modeling in zebrafish embryogenesis. DanioDecima was trained by transfer learning from Borzoi and Decima models trained on human and mouse genomes. We fine-tuned these pretrained transformer-based models on the Zebrahub single-cell RNA-seq atlas comprising 154 cell types across 10 developmental stages. Aggregating cells with the same cell type and developmental stage yielded 304 expression profiles (pseudobulks). Cell-type and timepoint-specific pseudobulks were generated following the protocol in Decima [7].

We used the 8-layer transformer architecture described in the Decima framework [7]. Model weights were initialized from checkpoints obtained from Human Decima, Human Borzoi, Mouse Borzoi, and through random initialization. Final model weights can be obtained at [https://doi.org/10.5281/zenodo.20348535]. Model inputs comprise 524,288 bp DNA sequences around each gene, and outputs are expression predictions per pseudobulk. We performed hyper-parameter optimization, focusing on learning rate and weight initialization strategies, evaluated using Pearson correlation and Poisson loss. Hyperparameter optimization identified learning rates of 3 × 10^−5^ for pretrained models and 3 × 10^−6^ for random initialization. Early stopping was implemented with validation loss differences less than 0.001 over 10 epochs. Weight decay tests did not improve performance significantly.

We inspected models by Pearson correlation, weighted Poisson loss, and multinomial loss. To avoid data leakage, dedicated chromosomes (chromosomes 3, 5, 14, 24, 25 for the validation set, and chromosomes 1, 4, 8, 9, 21 for the test set) are held out during training and used for evaluation. Integrated Gradients were used for sequence importance attribution analyses.

For regulatory element design, we adapted the evolutionary optimization workflow implemented in gReLU [18], starting from random sequence initialization. We compared designed sequences’ predicted tissue specificity and biological plausibility through motif enrichment analyses.

Code for the zebrafish adaptation, transfer learning experiments, and downstream analyses is available at https://github.com/czbiohub-sf/DanioDecima. A more detailed version of the methods can be found in the Supplementary Methods.

## 4 Experiments and Results

### 4.1 Transfer learning from mammalian sequence-to-expression models improves gene expression prediction in zebrafish embryogenesis

DanioDecima achieves robust predictive performance across diverse cell types, developmental stages and genes (Figure 2). Substantial performance improvement is observed when initializing models with pretrained weights compared to random. We observe an ≈ 0.20 increase in pseudobulk Pearson *r* and 0.14–0.15 in gene-level Pearson *r*.

**Figure 2:**
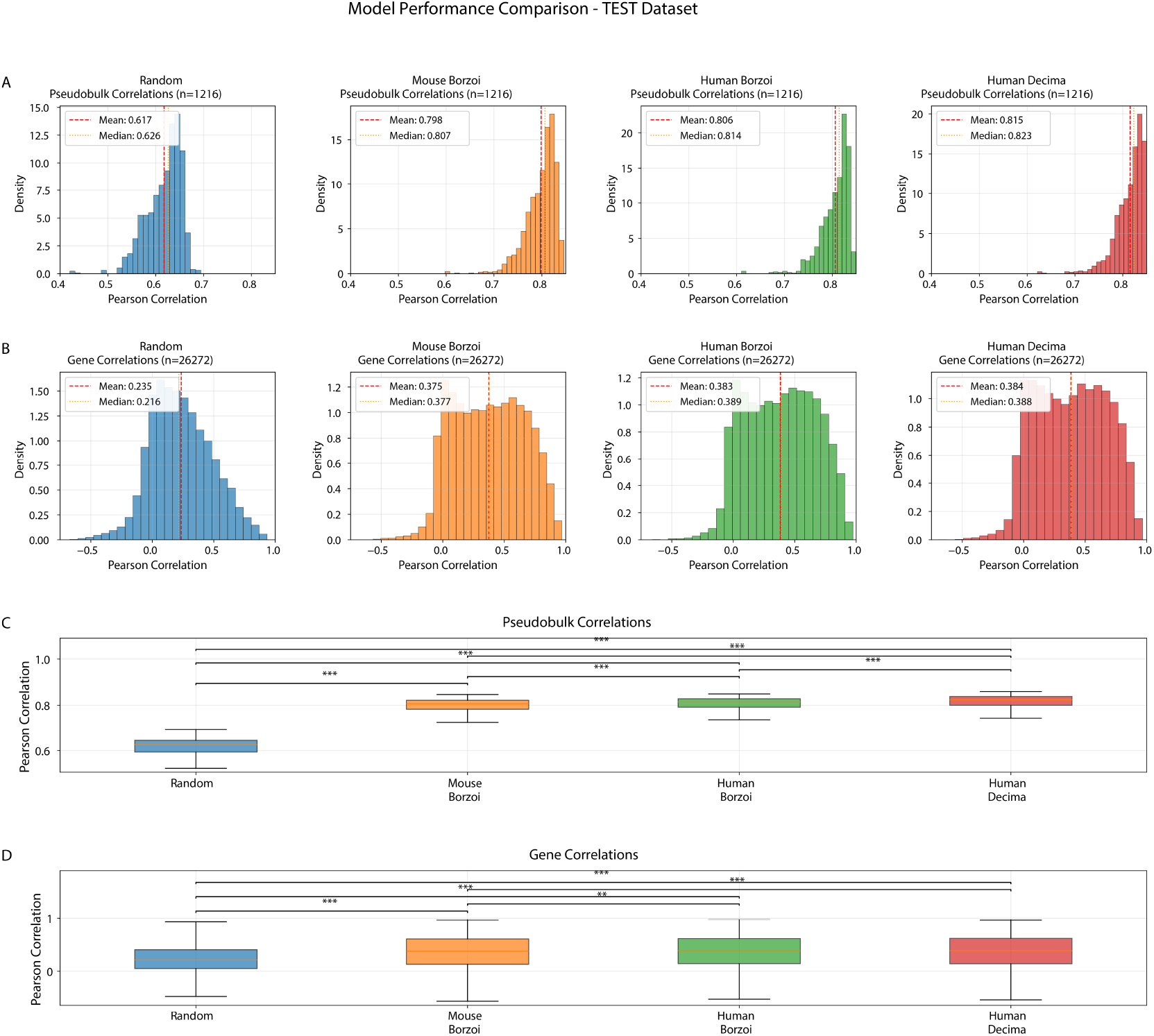
Predictive performance across zebrafish pseudobulks and genes improves with mammalian pretrained models. Transfer learning from mammalian sequence-to-expression models improves both cell-type–level and gene-level prediction accuracy relative to random initialization, with Human-Decima initializations yielding the strongest and most consistent performance gains. Upper panels: Histograms of Pearson correlations across test pseudobulks (n=1,216, 304 pseudobulks x 4 replicates). Red dashed lines mark the mean, yellow dotted lines indicate the median. Mean/median shifts from 0.62/0.63 (Random) to 0.81/0.83 (Human-Decima). Middle panels: Histograms for gene-level correlations across all test set genes. Mean/median rise from 0.24/0.22 to 0.38/0.39 in Human-Decima. Lower panels: Box plots comparing correlation distributions for pseudobulk (above) and gene-level (below), with pairwise statistical comparisons (Pairwise Mann-Whitney tests). Asterisks denote significance (* * *p <* 0.01, * * \**p <* 0.001).

Overall, performance improvements are statistically robust and widespread across thousands of pseudobulks and gene targets, reinforcing generalizable predictions. Notably, Human-Decima initialized models display the best performance, highlighting the benefit of initializing with weights of models pretrained with single cell functional genomics datasets over bulk tissue.

Transfer learning improves baseline performance most effectively in later embryonic stages, where the prediction task requires generalizing over heterogeneous cell types (Figure 9). The robust pattern across models and time points indicates that DanioDecima genuinely enhances sequence-to-expression prediction via learned regulatory priors, rather than superficial memorization. Notably, the best-performing pseudobulk predictions coincide with the mid-to-late phylotypic developmental stages (16-24 hpf) (Figure 9).

### 4.2 Transfer learning enhances predictive performance independent of gene conservation or expression level

To decouple how the possible memorization of gene element sequences contributes to expression level prediction, we analyzed predictive performance as a component of both expression level and gene conservation (Figure 3 A,D).

**Figure 3:**
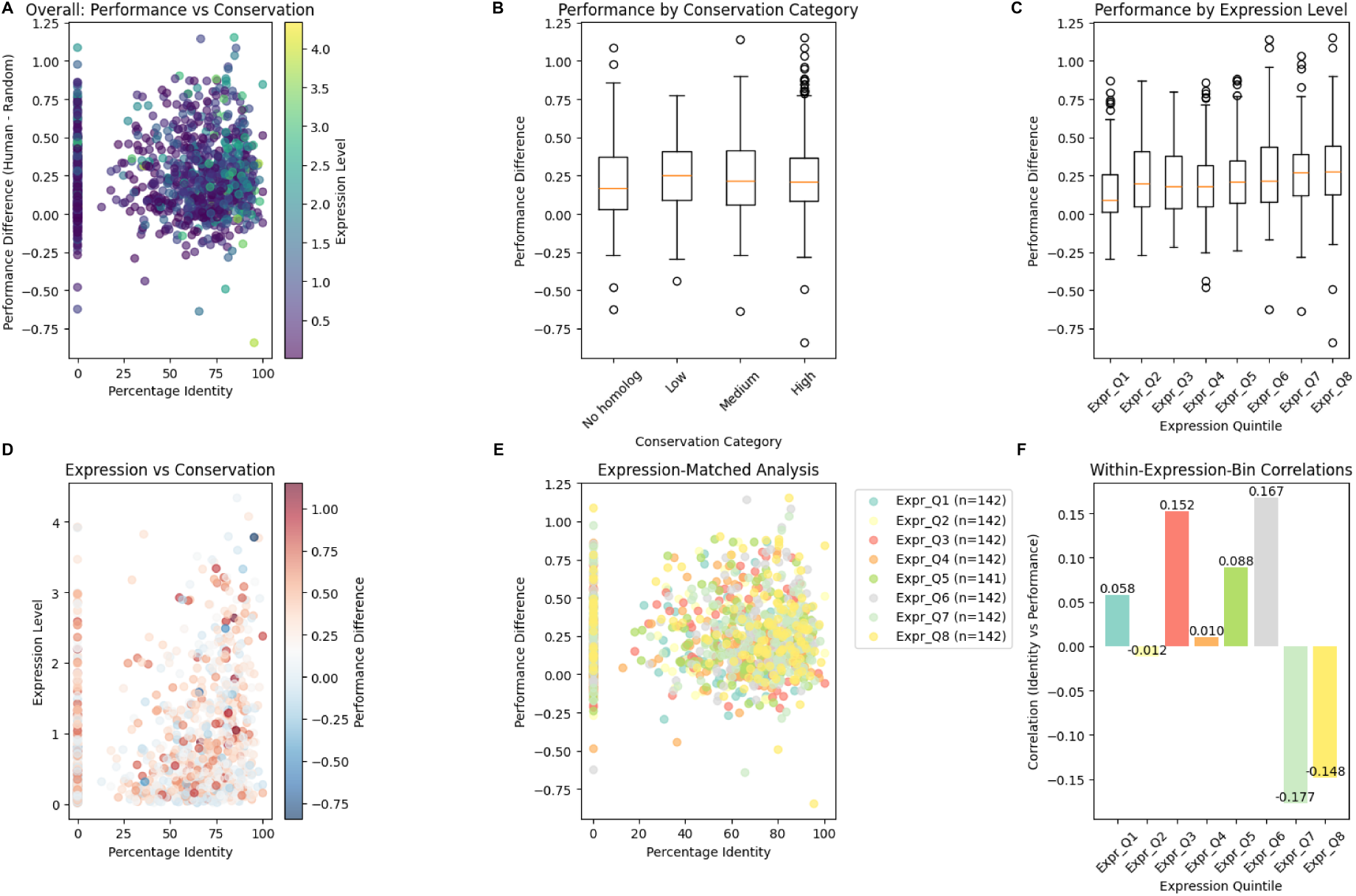
Transfer-learning gains show limited association with gene conservation and expression level. Performance improvement is defined as the per-gene Pearson *r* of the best Human-Decima-initialized model minus the random-initialization baseline. Each point is one of the ~1,135 held-out test genes. **A**, Improvement vs. percent sequence identity to the human ortholog, with each point colored by zebrafish expression level. No systematic trend is apparent; predictive improvement is broadly positive across the full conservation range, including genes with no detectable human homolog (left edge, ~0% identity). **B**, Same data binned into four conservation categories (No homolog, Low, Medium, High). Median improvement is comparable across categories (~0.18–0.25), so gains are not restricted to conserved orthologs. **C**, Predictive gains binned by expression level (octiles Q1–Q8, lowest to highest). Improvement is positive in every bin and rises modestly with expression (median ~0.10 in Q1 vs. ~0.27 in Q8). **D**, Joint distribution of expression and percent identity for each test gene, with point color encoding predictive improvement. No localized region of the (conservation, expression) plane drives the overall gain. **E**, Predictive improvement vs. percent identity replotted with color indicating expression octile, so that conservation effects can be inspected within expression-matched strata. The within-color spread is comparable to the across-color spread, indicating that conservation contributes little to predictive gains once expression is held approximately constant. **F**, Pearson correlation between percent identity and predictive improvement computed independently within each expression octile. All eight within-bin correlations are weak (|*r*| ≤ 0.18) and inconsistent in sign, confirming that, conditional on expression, sequence conservation explains little of the performance gain. Together, these panels show that transfer learning enhances regulatory inference broadly across genes, rather than through memorization of conserved promoter sequences.

A boost in performance is seen across the full conservation range and shows no systematic trend with sequence identity to the human ortholog (Figure 3 A,B). While the degree of improvement rises modestly with expression level (Figure 3 C), conservation explains little additional variance within fixed expression strata (within-octile |*r*| ≤ 0.18; Figure 3 E,F). Consistent gains in low-conservation genes show that the pretrained model is extracting regulatory grammar, rather than memorizing conserved promoter sequences.

### 4.3 Learned regulatory grammar corresponds to functional genome architecture

Attribution analyses (Figure 4 and Figure 5) confirm that pretrained models assign greater functional importance to genomic regions associated with biological regulation (promoters, introns, ATAC-seq peaks) compared to randomly initialized models.

**Figure 4:**
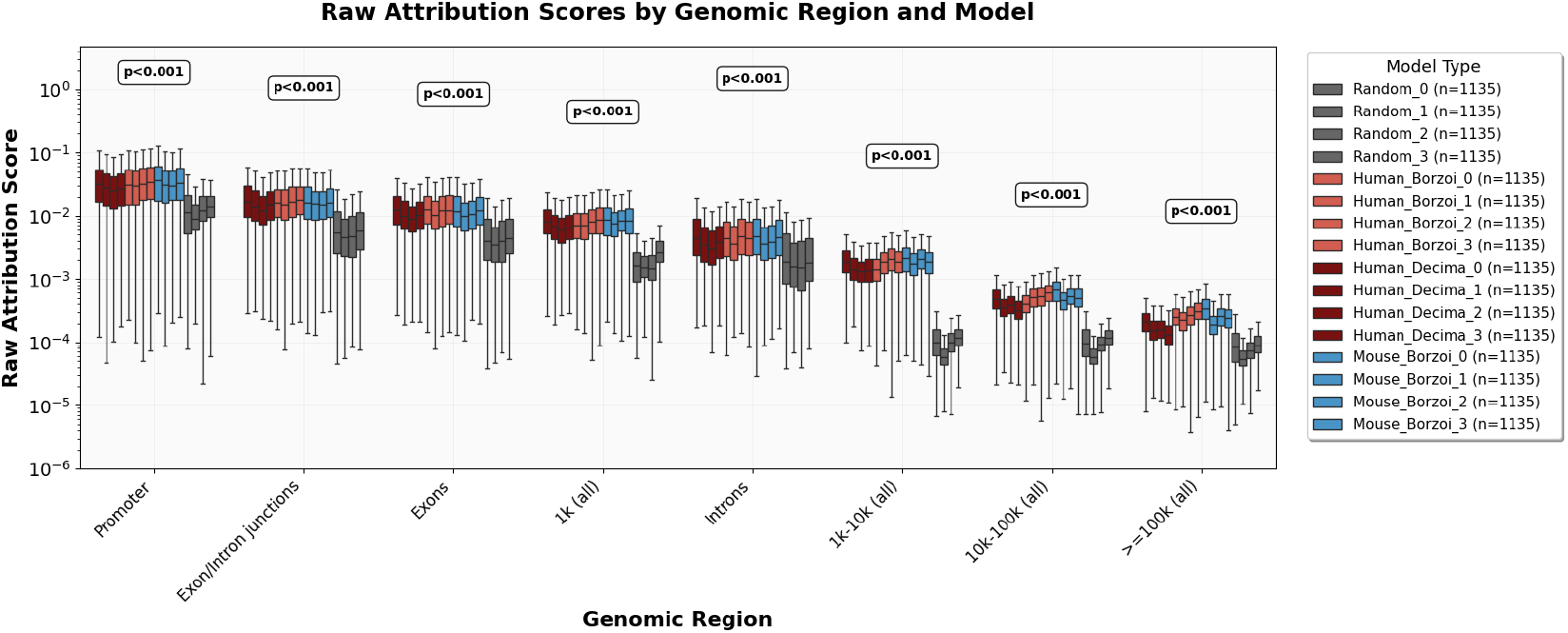
Pretrained models exhibit increased attribution to distal regulatory regions. Attribution patterns across genomic regions for different model initializations indicate that pretrained models place relatively greater importance on distal cis-regulatory elements alongside proximal promoter regions, consistent with known long-range regulatory interactions. Box plots show log-scale raw attribution scores (e.g. Integrated Gradients) aggregated across 1,135 test genes, comparing Random initialization (gray), Mouse-Borzoi (blue), Human-Borzoi (red), and Human-Decima (dark red). Pretrained models significantly emphasize promoter regions (p<0.001). Elevated attributions persist across distal intronic and intergenic bins (1–100kb and >100kb), demonstrating recognition of remote regulatory elements. All pretrained variants consistently outperform random baseline in assigning high importance to biologically relevant regulatory regions.

**Figure 5:**
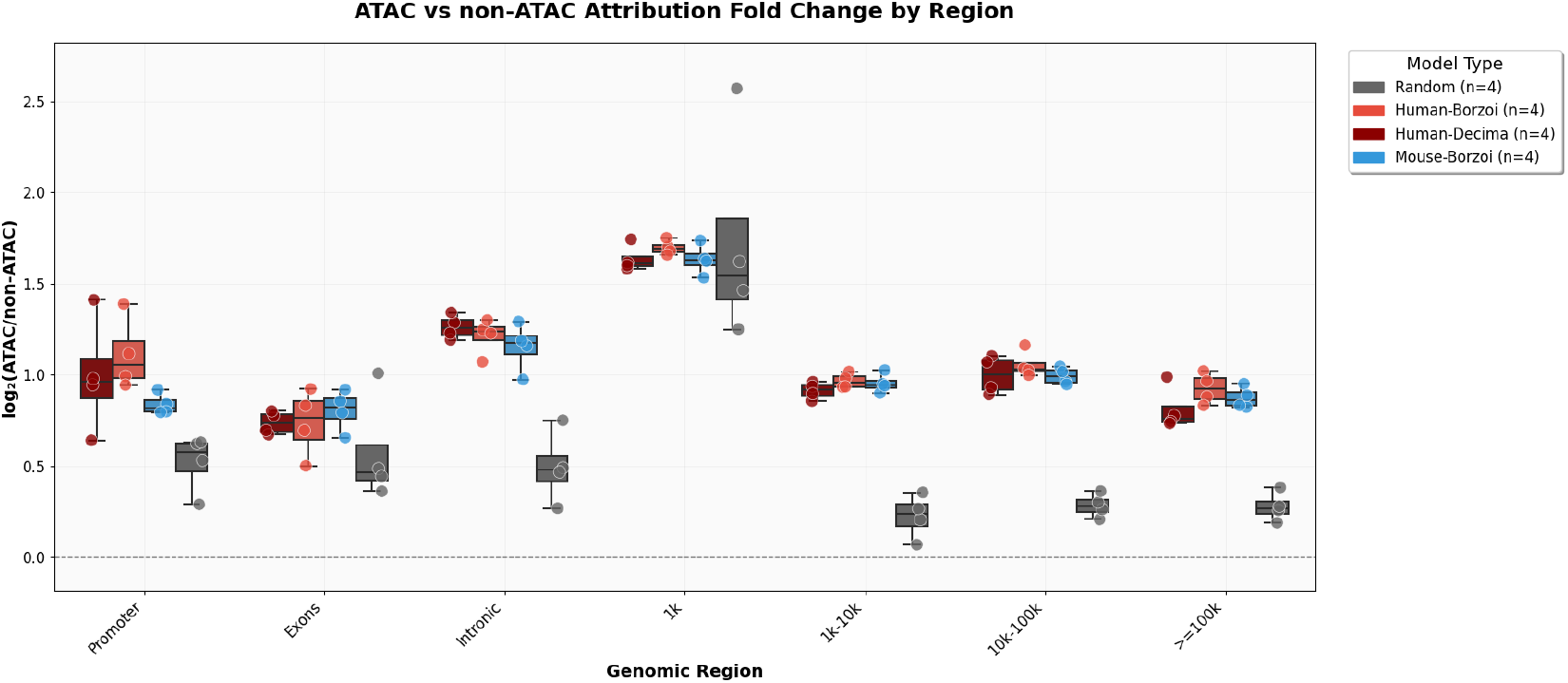
High attribution scores overlap with ATAC-seq peaks, highlighting biological relevance of the model’s predictions. DanioDecima and other pretrained models preferentially attribute importance to regions with biological evidence of accessibility and regulatory activity, demonstrating that the learned regulatory grammar corresponds to functional chromatin architecture as captured by ATAC-seq in zebrafish embryogenesis. The box plot shows the model attribution fold-change in ATAC compared to non-ATAC peak regions, stratified by genomic region and initialization. Each box plot represents the distribution of *log*_2_(ATAC / non-ATAC) attribution fold-change across 1,135 test genes for one of four initialization types: Random-initialized (gray), Mouse-Borzoi (blue), Human-Borzoi (light red), and Human-Decima (dark red). Pretrained models show substantial enrichment in attribution scores at ATAC-marked promoter and intronic regions, often 1.2–1.8× higher than non-ATAC controls (log fold-change ~0.3 −0.8). Moreover, randomly-initialized models show minimal enrichment, indicating poorer alignment with chromatin accessibility.

Models initialized with pretrained weights heavily emphasize regulatory regions and distal regions compared to random baselines (Figure 4). Attributions remain elevated across introns and distal bins up to >100kb. Randomly-initialized models, by contrast, show much lower scores beyond immediate promoter vicinity. Moreover, both Human-Decima and Borzoi variants yield similar attribution trends, outperforming random in all categories.

These observations align with regulatory biology: Enhancers and distal elements (often located tens to hundreds of kb from TSS) play significant roles in gene regulation via chromatin looping and TF binding (promoters alone are not sufficient). In addition, these results reflect model awareness of distal regulatory grammar, as the pretrained-model signal distribution across extended regulatory regions mirrors known cis-regulatory logic in developmental gene control.

ATAC-seq peaks mark active regulatory elements, such as promoters and enhancers, in the chromatin landscape (e.g. Pou5f3/Sox19b/Nanog controlled regions during zebrafish zygotic genome activation [19]). As attribution correlates with these peaks, it suggests the model is attuned to these regions as being important for transcript regulation, and not likely artifacts.

We find a significant overlap of model attribution with experimentally-verified ATAC-peaks compared to non-ATAC peak regions within relevant regulatory regions (Figure 5) (fold-change >1, typically 1.2–1.8). This suggests that pretrained models consistently assign more importance to ATAC-supported regions compared to randomly-initialized models.

### 4.4 DanioDecima enables the design of biologically plausible regulatory elements through directed evolution

Next we tested whether DanioDecima can be used to design regulatory elements that result in cell-type specific gene expression, a task that could allow for targeted interventions in the organism. Regulatory elements were designed using the suite of DanioDecima models through in-silico directed evolution as described in [7] (Fig. 6). Motif architectures designed for a given cell type (e.g. Heart, Neural, Mesoderm) consistently cluster together, indicating coherent motif enrichment strategies driven by the model (Fig. 6).

**Figure 6:**
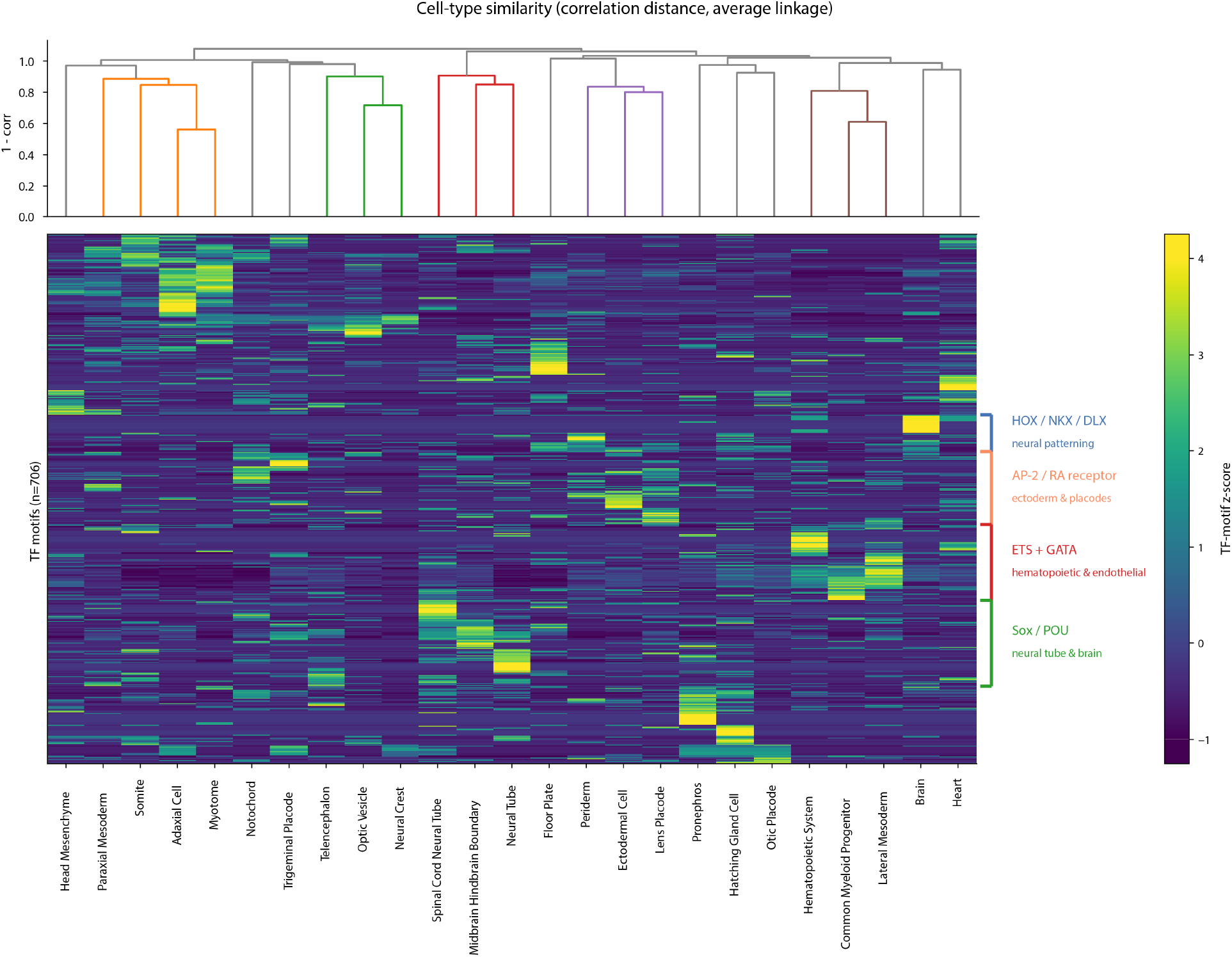
Bi-clustered motif-signature heatmap of DanioDecima-designed cell-type-specific regulatory sequences. Rows: 706 transcription-factor motifs identified by TF-MoDISco from the attribution scores of designed sequences. Columns: the 25 target cell types. Color: row-wise z-score of motif occurrence (viridis, z-score ∈ [−1, 4]; actual range [−1.42, 4.80] clipped to the rectangular colorbar caps). Columns are clustered by correlation distance with average linkage (top dendrogram; branch heights = 1 −Pearson correlation; colored sub-trees mark clusters detected at 85% of the maximum linkage distance). Rows are clustered by Euclidean distance with average linkage (row dendrogram omitted for legibility). Four motif clusters with the strongest TF-family signature are bracketed on the right: HOX / NKX / DLX (neural patterning), AP-2 / RA receptor (ectoderm and placodes), ETS + GATA (hematopoietic and endothelial), and Sox / POU (neural tube and brain). Cell types from related lineages cluster together, and TF-motif modules co-vary block-diagonally with these cell-type clusters, demonstrating that DanioDecima learns lineage-specific regulatory grammars and supporting its use as a design oracle for cell-type-specific regulatory elements.

These patterns support the notion that DanioDecima’s in silico evolutionary design reliably encodes distinct regulatory programs per cell type, which align with known developmental transcription factor usage, such as GATA motifs in endoderm, SOX/POU in neural lineages, etc., consistent with zebrafish chromatin accessibility and motif enrichment datasets like Zebrahub and scATAC-seq atlases [15, 20].

## 5 Discussion

DanioDecima demonstrates the effective transfer of regulatory knowledge from mammalian systems to zebrafish in sequence to function modeling, despite approximately 450 million years of evolutionary divergence. The model captures fundamental regulatory logic conserved across vertebrates, yet also adapts to zebrafish-specific regulatory patterns.

Beyond predictive performance, the regulatory element design pipeline illustrates how DanioDecima can practically inform synthetic biology and developmental genetics experiments. This work also highlights the value of community resources such as gReLU [18], which facilitates reproducible benchmarking, model developments and sequence design across regulatory genomics tasks.

The strong performance of transfer learning in DanioDecima is consistent with emerging evidence from plant regulatory modeling [21] and cross-species scRNA-seq integration [22] that deep models can learn biological representations that transfer across species boundaries. Together, these findings support the broader hypothesis that many regulatory principles are evolutionarily conserved and therefore amenable to shared representation learning approaches.

While DanioDecima demonstrates gains in predictive performance, several caveats should be considered. Deep sequence models trained across species can overestimate cross-species generalization if orthologous genomic regions inadvertently appear in both training and testing splits. Recent work has shown that such leakage can inflate performance metrics like Pearson *r*, making disentangling memorization from true regulatory inference challenging [23]. In this study, we address this concern by investigating the correlation between a gene’s conservation and predicted performance gain. However, future work could implement strict orthology- or chromosome-based holdouts to remove such biases. In addition, the use of pseudobulked scRNA-seq data may obscure biologically important single-cell heterogeneity. Aggregating scRNA-seq data into pseudobulks risks obscuring rare cell states, stage-specific noise profiles, and ambient RNA contamination. These factors are known to affect downstream modeling. Dedicated single-cell models or sub-sampling schemes could complement bulk metrics.

Several directions exist for future work. Experimental validation using in vivo reporter assays or CRISPR-based perturbation systems will be important for confirming the functional activity of designed regulatory elements and establishing the practical utility of the framework. Incorporating additional modalities including spatial transcriptomics, chromatin accessibility, and imaging data may further improve predictive resolution and enable richer modeling of developmental processes. In particular, incorporating multiomic constraints and modeling cellular trajectories through time with autoregressive or diffusion-based architectures may improve resolution and support applications in temporal gene regulation and lineage specification.

Further investigation is needed to define the limits of cross-species transferability. Systematic evaluation across organisms with varying evolutionary distances, genome architectures, and regulatory mechanisms will help clarify which components of regulatory grammar are broadly conserved versus species-specific. In parallel, further exploration into alternative initialization strategies using large DNA language models such as Evo 2 [9] and related foundation models may further improve biological generalization in cross-species regulatory modeling.

## 6 Acknowledgements

We thank the Computational Biology Platform and the Royer group of the Biohub for helpful discussion and support. We thank the Scientific Computing team for their support for HPC. We appreciate the brainstorming and feedback from Reinier Grona, Sayan Ghosal, and Giovanni Palla. Funding for this work was provided by Biohub. We thank the Biohub donors, Priscilla Chan and Mark Zuckerberg, for their generous support.

## A Supplementary Results

### A.1 Model Training

Figure 7 shows the validation loss, training loss, and Pearson *r* correlations over epochs of training. Models initialized with pretrained weights (Human-Decima and Borzoi) outperform random initializations. Specifically, Human-Decima-initialized models reach higher validation Pearson correlations earlier (*≈* 0.82), compared to Borzoi (*≈* 0.80) and random initializations (*≈* 0.60).

**Figure 7:**
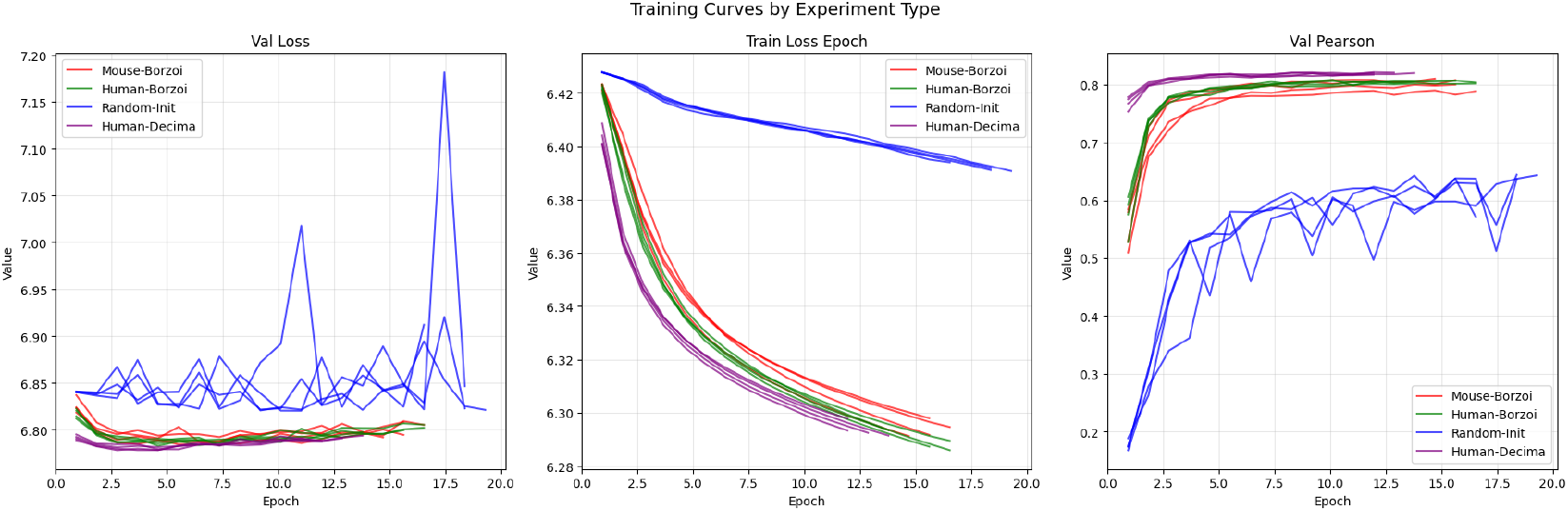
Validation loss, training loss, and Pearson *r* curves over epochs of model training. Initializing with pretrained model weights significantly boosts predictive performance, with Human-Decima pretrained weights outperforming Borzoi and random initializations. Transfer learning from mammalian pretraining substantially enhances predictive accuracy in zebrafish embryogenesis models.

A learning rate sweep (Figure 8) confirmed optimal learning rates at 3 × 10^−5^ for models initialized with pretrained weights, and 3 × 10^−6^ for randomly initialized models.

**Figure 8:**
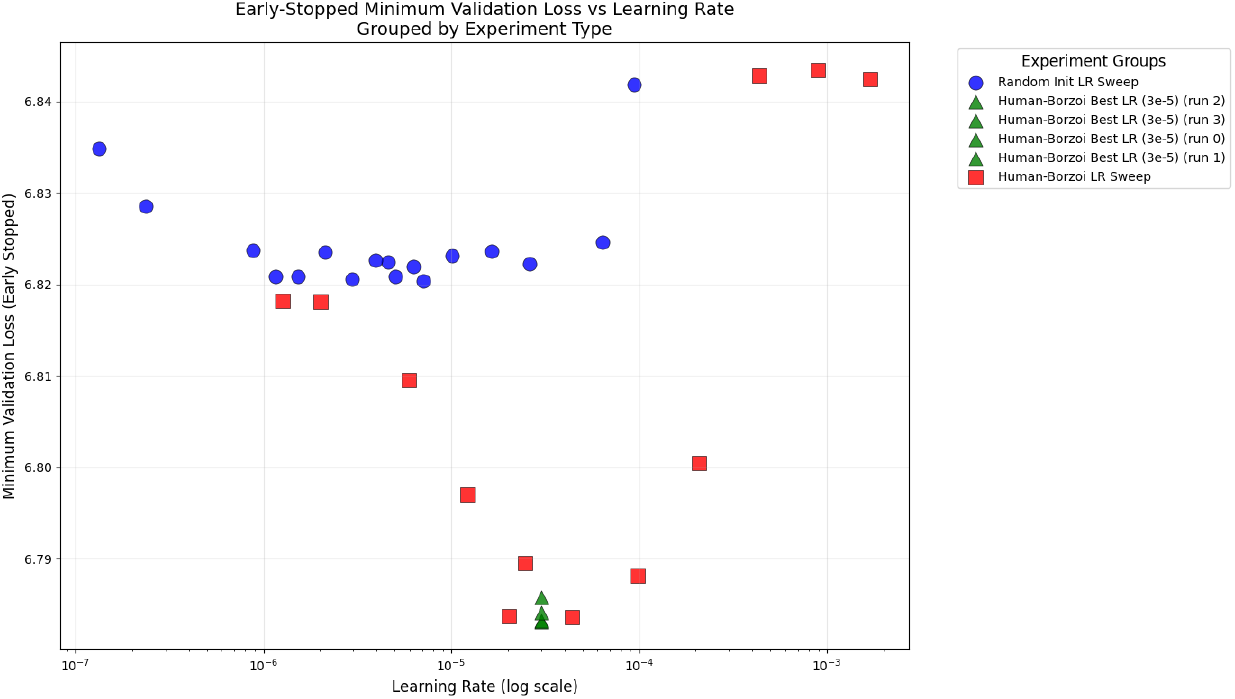
A learning rate sweep demonstrates that Human-Borzoi pretraining achieves the lowest early-stopped validation loss at a learning rate of 3 ×10^−5^, matching the optimal setting for Human-Decima [7] Randomly initialized models reach best performance at a lower learning rate of 3 × 10^−6^, suggesting that pretraining enables more stable and effective optimization at higher learning rates.

### A.2 Predictive Performance Across Cell Types and Developmental Stages

The best-performing pseudobulk predictions coincide with the mid-to-late phylotypic developmental stages (16-24hpf) (Figure 9). To diagnose this effect more precisely, we assess model performance improvements over performance of the randomly-initialized models at the given time points (Figure 10). The largest relative improvements are observed from 16hpf to 3dpf, which may reveal consistencies with the “hourglass” model in developmental biology. Random-initialized models show more poor performers in late organogenesis (3–10 dpf), suggesting less regulatory predictability post-phylotypic period, where pretrained models still maintain higher relative performance.

**Figure 9:**
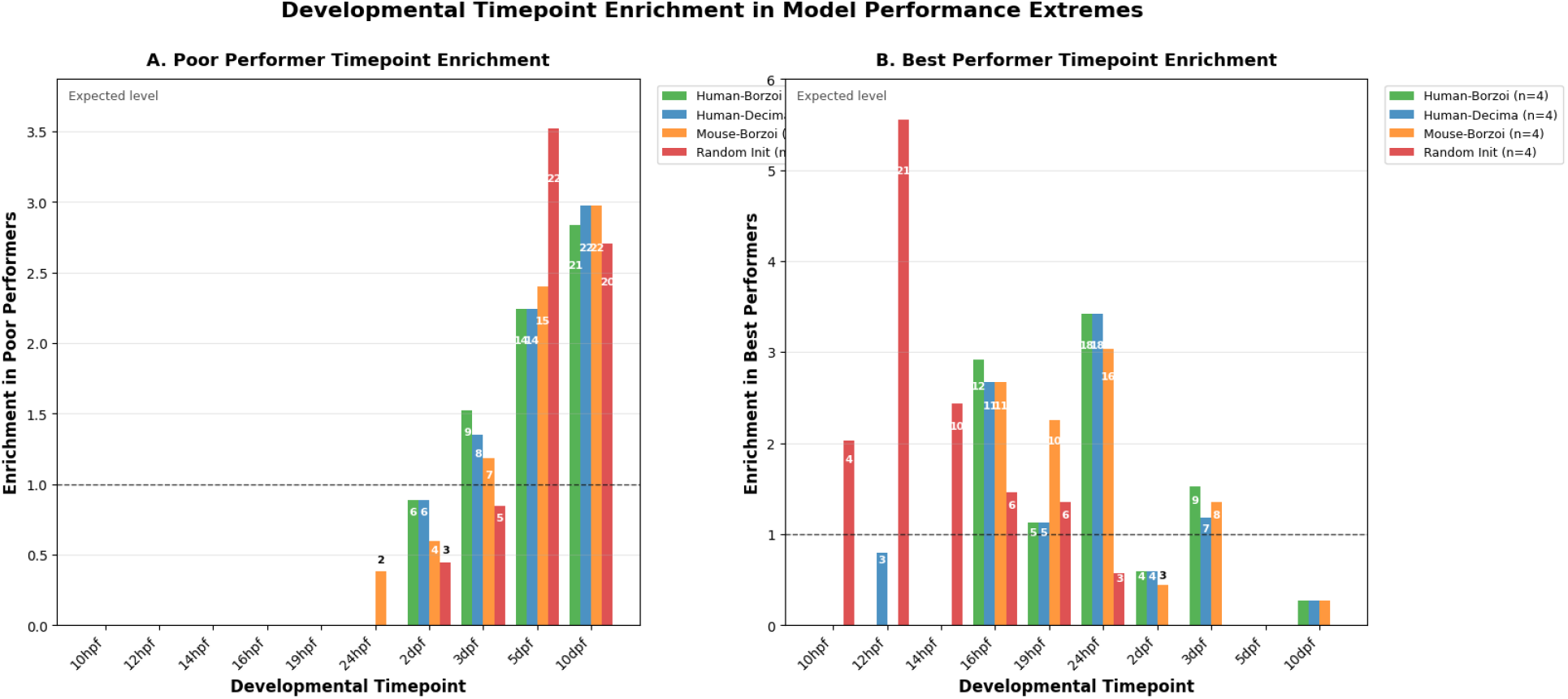
Distribution of best- and worst-performing pseudobulks across developmental stages. Developmental-stage enrichment of best and worst performing pseudobulks across pretrained and randomly initialized models **A**, Enrichment of the lowest-performing (bottom 4) pseudobulk samples over developmental time points for each initialization strategy. Poorer performance is found in later stages for all model initialization strategies. **B**, Enrichment of the highest-performing (top 4) pseudobulk samples by time point. Compared to randomly initialized models, top performers consistently cluster around the phylotypic window (16–24 hpf) across pretrained strategies, indicating regulatory predictability is highest in this mid-developmental stage.

**Figure 10:**
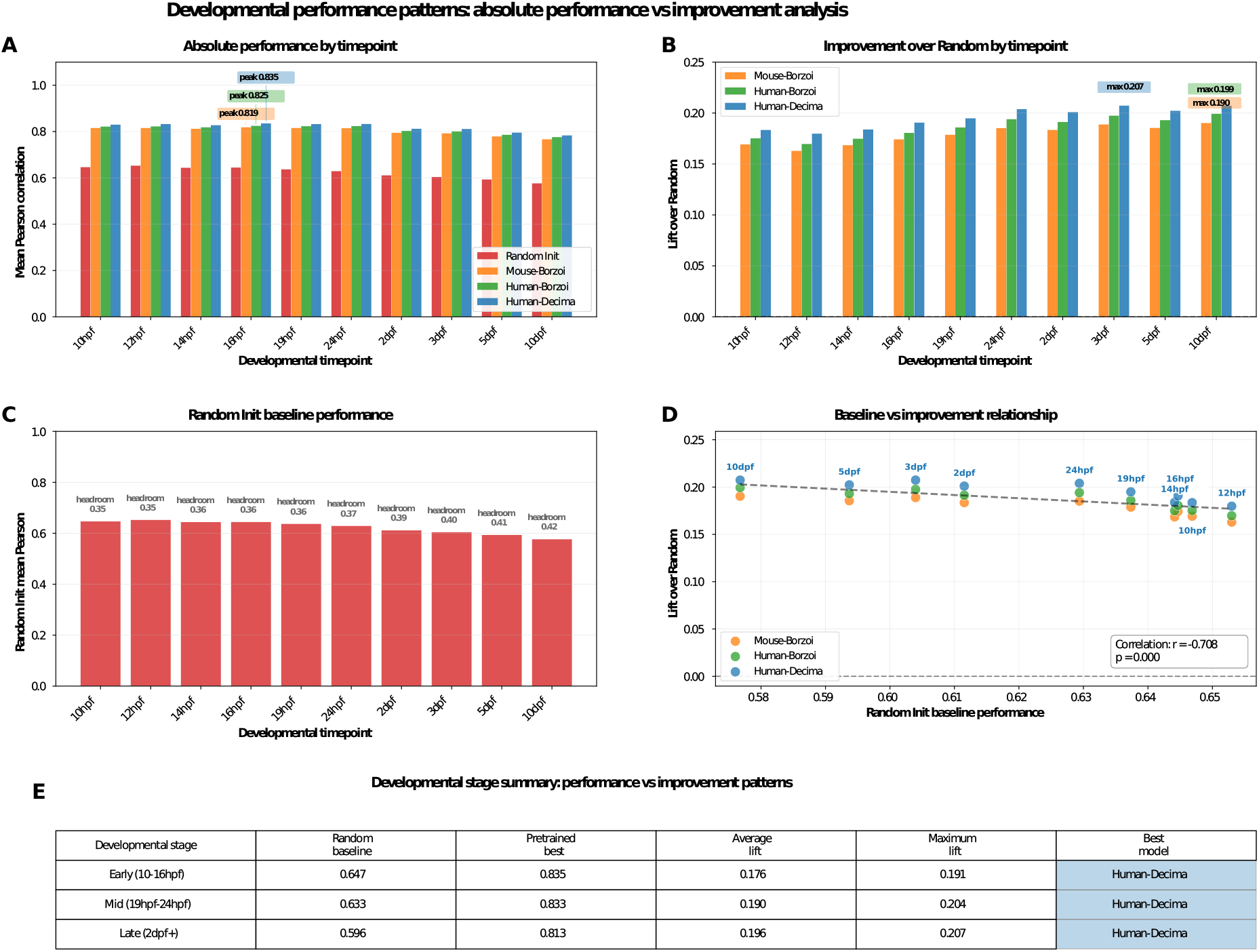
Developmental-stage specific predictive performance and improvement over baseline. Performance diagnostics highlight greatest improvements at later stages (2dpf+), due to lower baseline performance of random models. **A**, Pearson correlation between predicted and actual gene expression for randomly initialized, Mouse-Borzoi–initialized, Human-Borzoi, and Human-Decima, plotted across 10 embryonic timepoints (10hpf to 10dpf). Human-Decima achieves highest absolute performance, peaking at 0.84 at 16hpf. **B**, Improvement in Pearson r over random initialization for each pretraining strategy. Human-Decima consistently achieves the greatest improvement, with a maximum of 0.21 at 3dpf. **C**, Baseline performance of the random model per timepoint, showing a decline from 0.65 in early development to 0.58 by 10dpf. **D**, Scatter plot correlating baseline random performance (x-axis) with improvement (y-axis) across timepoints and models. There is a strong negative correlation (*r ≈ −* 0.71, *p*′0.001), indicating larger improvement when the baseline is weaker. Some time points show higher improvements than expected (distance from linear correlation dashed line). **E**, Summary table aggregates stages (Early 10–16hpf; Mid 19–24hpf; Late 2dpf+), showing average and maximum improvement values; Human-Decima is the best performing model in all windows.

## B Supplementary Methods

We built upon the gReLU framework [18] and the Decima modeling pipeline [7], which provide infrastructure for genomic sequence modeling, training, interpretation, and regulatory sequence optimization workflows. Our work extends these frameworks to cross-species transfer learning and regulatory modeling in zebrafish embryogenesis. Unless otherwise specified, implementation details, optimization procedures, and attribution methods follow the original framework implementations.

### B.1 Data Exploration and Preprocessing

Dataset characteristics were analyzed through comprehensive exploration of zebrafish single-cell RNA-seq data [15] aggregated into pseudobulk profiles following the protocol from [7]. The analysis examined gene expression variance patterns to identify highly variable genes using scanpy’s variance filtering approach (min_mean=0.0125, max_mean=5, min_disp=0.5). This filtering identified genes most likely to benefit from improved predictive modeling, with highly variable genes showing stronger correlations between gene variance and model performance differences compared to uniformly expressed genes.

### B.2 Model Training

DanioDecima models were fine-tuned using an experimental design with 16 total configurations across four initialization strategies:

1. **Random Initialization (n=4):** Models initialized with random weights using different seeds and learning rate 3×10^−6^.
2. **Pretrained Human-Borzoi (n=4):** Models initialized from human Borzoi checkpoints with learning rate 3×10^−5^.
3. **Pretrained Mouse-Borzoi (n=4):** Models initialized from mouse Borzoi checkpoints with learning rate 3×10^−5^.
4. **Pretrained Human-Decima (n=4):** Models initialized from human Decima checkpoints with learning rate 3×10^−5^.

Training employed SLURM array jobs with GPU allocation, using PyTorch Lightning for distributed training. Each model was trained for up to 40 epochs with early stopping (patience=10, min_delta=0.0001) and gradient accumulation (batch_size=4, accumulate_grad_batches=5). Models were trained using the TaskWisePoissonMultinomialLoss implemented in the Decima framework and optimized with Adam using configurable weight decay.

### B.3 Generating Model Predictions

Model predictions were generated using the Decima inference pipeline applied across all trained checkpoints. Predictions included:

- Batch prediction across all pseudobulk profiles
- Computation of per-gene Pearson correlations between observed and predicted expression
- Per-dataset pseudobulk correlation calculations
- Integration of predictions into AnnData format with metadata

### B.4 Model Performance Evaluation

#### B.4.1 Celltype-Specific Analysis

Performance evaluation focused on cell-type-specific metrics from the evaluation notebook `daniodecima-applications-main/notebooks/4_evaluation/01_evaluate_celltypes.ipynb`. The analysis examined:

- **Pseudobulk Correlations:** Mean Pearson correlations calculated across developmental timepoints for each cell type, providing tissue-specific performance measures.
- **Gene-Level Performance:** Per-gene correlations aggregated across cell types, identifying genes most amenable to improved prediction.
- **Developmental Stage Analysis:** Performance stratified by embryonic timepoints (10hpf through 10dpf), revealing temporal patterns in model effectiveness.
- **Statistical Comparisons:** Pairwise Mann-Whitney U tests with FDR correction comparing model performance across cell types and developmental stages.

#### B.4.2 Performance Metrics

Primary evaluation metrics included mean Pearson correlation for both pseudobulk profiles and individual genes, with additional analysis of performance variance, percentile ranks, and improvement magnitudes relative to random initialization baseline.

### B.5 Attribution Analysis for Conservation Confounding

This workflow examines model interpretability through attribution analysis to assess potential confounding between model predictions and evolutionary conservation patterns.

#### B.5.1 Attribution Score Calculation

Attribution analysis was performed using the combined attribution calculation pipeline (`daniodecima-applications-main/notebooks/5_specificity/00_combined_attribution_analysis.py`) submitted via SLURM array jobs across all 16 experimental model configurations. Using the attribution utilities provided in the Decima and gReLU frameworks, we performed the following analyses:

- **Model Loading:** Each trained DanioDecima checkpoint was loaded and configured for inference mode with gradient computation disabled for computational efficiency
- **Attribution Method:** InputXGradient attribution, as implemented through Captum and integrated into the Decima interpretation pipeline, was used to calculate nucleotide-level importance scores.
- **Task Aggregation:** Cell-type specific attributions were computed using task-wise aggregation with mean pooling across target cell types (threshold *>* 0.5 expression)
- **Genomic Integration:** Attributions were computed across full 524,288 bp genomic windows and aggregated across nucleotide channels for downstream analyses.

The attribution pipeline processes each gene individually, computing attributions for cell types with detectable expression levels and aggregating results across the first four sequence channels (A, T, G, C) while excluding the gene mask channel.

#### B.5.2 Genomic Feature Annotation

Attribution scores were analyzed across distinct genomic regions to assess model focus patterns:

- **Gene Structure:** Promoters (*±*100bp from TSS), exons, introns, and exon-intron junctions (*±*10bp) were annotated using GTF coordinates
- **Regulatory Elements:** ATAC-seq peaks from zebrafish multi-omics data [20] were overlapped with attributions to identify accessible chromatin regions
- **Distance Categories:** Attributions were stratified by distance from gene bodies: 1kb, 1–10kb, 10–100kb, and *>*100kb flanking regions
- **Region-Specific Analysis:** Mean attribution scores were calculated separately for CRE (ATAC peak) and non-CRE regions within each genomic category

#### B.5.3 Conservation Confounding Analysis

This analysis examines whether high-performing models exhibit attribution patterns that correlate with evolutionary conservation rather than cell-type specific regulatory signals:

- **Gene Conservation Metrics:** Human-zebrafish ortholog conservation scores were integrated with model attribution patterns
- **Expression Controls:** Gene expression level distributions were used as covariates to control for expression-dependent attribution biases
- **Statistical Framework:** Mann-Whitney U tests with FDR correction were applied to assess differential attribution patterns between conserved and non-conserved genomic regions
- **Propensity Score Matching:** Expression-level matched analyses were performed to isolate conservation-specific effects from general expression-level confounding

### B.6 Regulatory Element Design with Cell-Type Specificity

We adapted the directed evolution framework introduced in Decima [7] and gReLU [18] to design synthetic zebrafish regulatory elements with enhanced cell-type specificity.

#### B.6.1 Evolutionary Design Framework

Regulatory element optimization was performed using the Decima sequence evolution pipeline with the following details:

- **Target Definition:** Cell-type specificity was defined as the difference between mean predicted expression in target cell types versus background cell types at 16hpf developmental timepoint
- **Sequence Initialization:** 200 bp promoter elements were initialized with random sequences and placed upstream of EBFP reporter cargo sequences
- **Genomic Context:** Following the convention established in the Decima evolution protocol [7], we placed the synthetic 200 bp regulatory element at zebrafish chromosome 4 position 29,480,218 (GRCz11), offset 163,840bp into the 524,288bp model input window. This position is intergenic - the nearest annotated protein-coding gene (*znf1140*) is approximately 52 kb away - so the placement avoids overwriting an endogenous regulatory element and lets the model’s predictions depend on the inserted sequence rather than on any nearby promoter. The choice is conventional rather than biologically motivated for zebrafish; cell-type specificity is enforced by the prediction objective (target - background mean), not by the insertion locus.

#### B.6.2 Directed Evolution Algorithm

Following the Decima evolution protocol, iterative single-point mutagenesis was used to optimize regulatory sequences:

- **Mutation Space:** All possible single nucleotide substitutions (3 per position × sequence length) were evaluated at each evolutionary round
- **Fitness Function:** Specificity scores calculated as mean target cell-type expression minus mean background cell-type expression from model predictions
- **Selection Strategy:** The highest-scoring mutation was selected and applied to the sequence each round, with tracking of position-specific evolution patterns
- **Convergence Criteria:** Early stopping implemented with patience=50 rounds and minimum improvement threshold=0.001 to detect evolutionary plateau

#### B.6.3 Sequence Analysis and Interpretation

Final evolved sequences were further characterized using in silico mutagenesis and transcription factor motif analyses provided through the Decima interpretation workflow:

##### In Silico Mutagenesis (ISM)

- **Systematic Perturbation:** Every position in final evolved sequences was mutated to all alternative bases (3 mutations per position)
- **Effect Quantification:** Prediction changes calculated relative to original sequence across all cell types
- **Visualization:** ISM effects displayed as position-specific line plots and base-substitution heatmaps to identify critical regulatory positions

##### Transcription Factor Motif Analysis

- **Motif Scanning:** JASPAR2020 vertebrate transcription factor binding site database used for comprehensive motif detection
- **ISM Weight Integration:** Motif hits weighted by underlying ISM attribution scores to prioritize functionally important binding sites
- **Regulatory Prediction:** High ISM-weight motifs identified as candidate drivers of cell-type specificity

#### B.6.4 Comparative Analysis and Visualization

##### Motif Clustering

Hierarchical clustering analysis revealed regulatory element design patterns:

- **Distance Metrics:** Correlation-based clustering of cell-type expression profiles using average-linkage
- **Heatmap Visualization:** Z-score normalized expression patterns displayed with dendrograms showing cell-type similarity relationships
- **Pattern Recognition:** Cluster analysis identified groups of cell types with shared regulatory sequence requirements

